# A Meiotic B-type Cyclin Selectively Drives Mitotic Progression and Fidelity in Cancer Cells

**DOI:** 10.64898/2026.09.28.754947

**Authors:** Amrutha Kizhedathu, Anh Cao Ngoc Nguyen, Clinton Yu, Shabnam Moghareh, Vy Tran, Powel Mousaian, Johnny Vertiz, Claudia A. Benavente, Lan Huang, Pablo Lara-Gonzalez

## Abstract

Cell cycle transitions are driven by cyclins in complex with cyclin-dependent kinases (CDK). Here, we demonstrate that cyclin B3, an evolutionarily divergent B-type cyclin previously thought to function exclusively in female meiosis, is expressed in mitotic cells and interacts with both CDK1 and CDK2. We show that cyclin B3-CDK1/2 is essential for timely mitotic progression by ensuring proper chromosome alignment and stable kinetochore-microtubule interactions, while its overexpression accelerates mitosis and improves chromosome alignment kinetics. Through phospho-mass spectrometry, we identify several kinetochore and mitotic spindle components as substrates of cyclin B3-CDK1/2 in mitosis. Notably, cyclin B3 depletion disrupts mitotic progression in most cancer cell lines tested – including colorectal, osteosarcoma, and breast cancer models – while non-transformed cells are largely unaffected. These findings identify cyclin B3-CDK1/2 as a novel driver of mitotic progression and reveal a preferential dependency in a subset of cancer cells that may represent a potential therapeutic vulnerability.

## INTRODUCTION

Progress through the cell division cycle is ensured by cyclin-dependent kinases (CDKs), which partner with proteins known as cyclins to phosphorylate a wide range of cellular substrates in order to facilitate cell cycle events (Pellarin et al., 2025; Tatum and Endicott, 2020; Wood and Endicott, 2018). Despite more than five decades of cell cycle research, how different cyclin-CDK complexes promote cell cycle transitions remains incompletely understood. This question is particularly relevant in cancer, where disrupted cell cycle regulation can fundamentally alter cell cycle dynamics.

The mitosis phase of the cell cycle is driven primarily by cyclin B in complex with CDK1 (Crncec and Hochegger, 2019; Lindqvist et al., 2009). The cyclin B family comprises three members: cyclin B1, cyclin B2, and cyclin B3 (Tatum and Endicott, 2020). While cyclin B1 and B2 share high sequence similarity, cyclin B3 is evolutionarily divergent and shares partial sequence homology with A-type cyclins (Gallant and Nigg, 1994; Lozano et al., 2002; Nguyen et al., 2002; Tschop et al., 2006). It is widely assumed that in vertebrates, cyclin B1 and cyclin B2 are the main mitotic B-type cyclins (Crncec and Hochegger, 2019; Lindqvist et al., 2009), while cyclin B3 roles are specific to female meiosis, where it promotes the metaphase-to-anaphase transition during meiosis I (Karasu et al., 2019; Li et al., 2019; Zhang et al., 2015). Indeed, female patients carrying homozygous loss-of-function mutations in cyclin B3 experience miscarriages that are attributed to oocyte maturation defects but are otherwise healthy (Fatemi et al., 2021; Rezaei et al., 2022; Wang et al., 2023).

Here, we unexpectedly find that cyclin B3 is expressed during mitosis, where it is critical for mitotic progression and fidelity by ensuring efficient kinetochore-microtubule attachments. Strikingly, we show that this function is conserved in a broad range of cancer cell types, while largely dispensable in non-transformed cells. Moreover, we find that cyclin B3-CDK1/2 drives the phosphorylation of hundreds of mitotic substrates involved in processes such as kinetochore function and spindle assembly. We further show that cyclin B3 overexpression is sufficient to accelerate mitosis and enhance its accuracy. Our findings reveal a surprising role for cyclin B3-CDK1/2 in cancer mitosis to support timely and accurate mitotic progression, which points to a potential vulnerability in cancer cell division that may be exploited for therapeutics.

## RESULTS AND DISCUSSION

### Cyclin B3 is expressed in mitosis and interacts with CDK1 and CDK2

To test if cyclin B3 is expressed in non-meiotic tissues at the protein level, we generated a polyclonal anti-cyclin B3 antibody (**Fig. 1A**). Strikingly, immunoblotting using this antibody revealed the presence of a specific band at the expected ∼158 kDa size for cyclin B3, whose levels were higher in mitotic HeLa cells compared to asynchronous extracts (**Fig. 1B&C**), indicating cyclin B3 expression in mitotic cells. Synchronous cell cycle extracts revealed that endogenous cyclin B3 begins to accumulate in late S-phase and early G2 (**Fig. 1D**); an accumulation profile that fits the pattern of a mitotic B-type cyclin. To define the CDK binding partner of cyclin B3 in mitosis, we performed immunoprecipitation analyses from HeLa cells stably expressing 3xFlag tagged cyclin B3 (**Fig. 1E**). We found that cyclin B3 co-immunoprecipitated with both, CDK1 and CDK2 (**Fig. 1E**; **S1A**). Using this cell line, we also found that cyclin B3 displayed a diffuse localization pattern during mitosis (**Fig. S1B**). Thus, unlike cyclin B1, cyclin B3 can efficiently partner with both CDK1 and CDK2 in mitosis.

**Figure 1.**
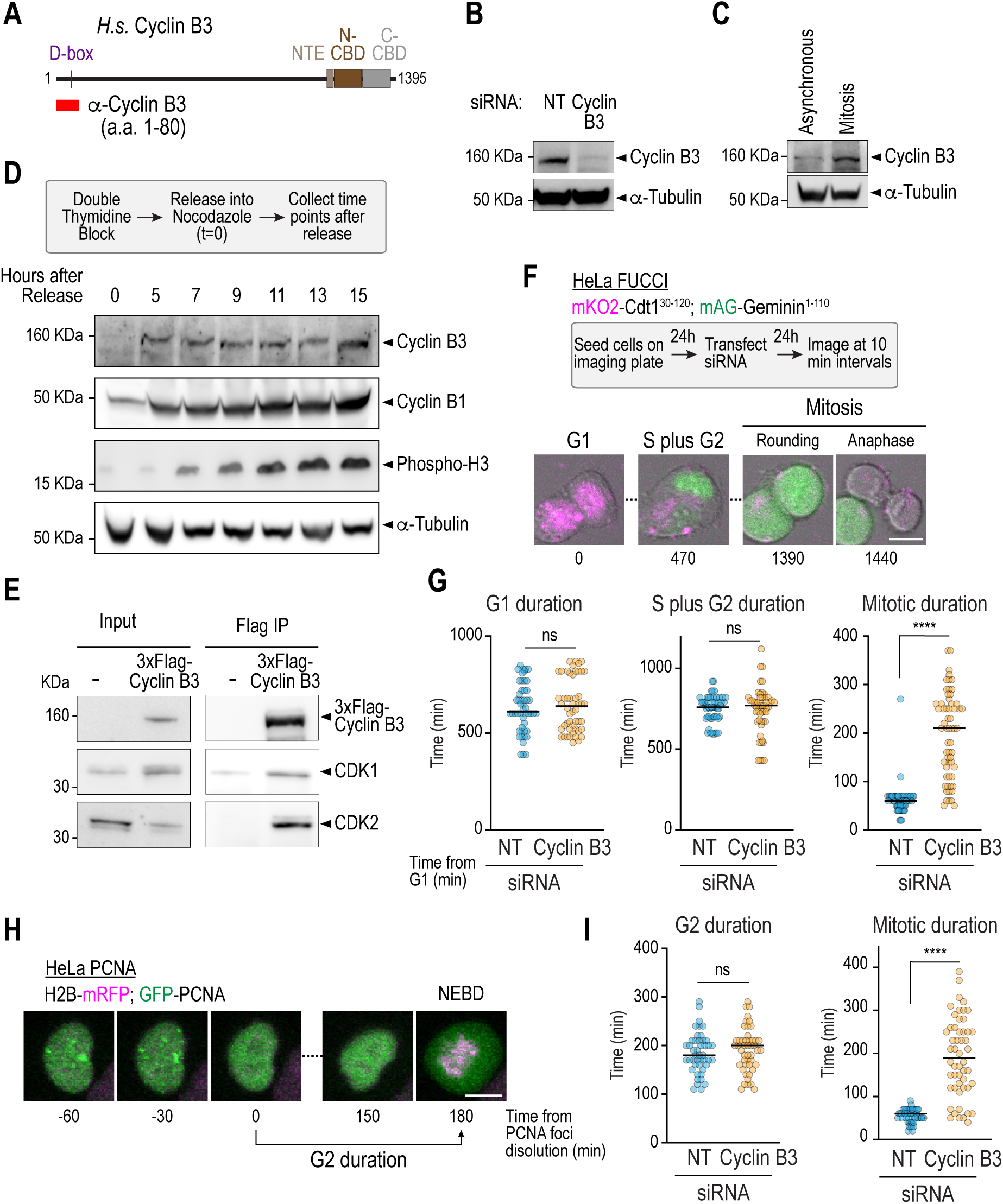
Cyclin B3 is expressed in mitosis and interacts with CDK1 and CDK2. **(A)** Schematic illustrating the structure of human cyclin B3. Key predicted domains in the protein (D-box, N-terminal extension (NTE) and Cyclin Box Domains (CBD)) are shown. The region used to raised the anti-cyclin B3 antibody (amino acids 1-80) is illustrated in red. **(B)**&**(C)** Immunoblots of HeLa cell extracts, treated with non-targeting siRNA (NT), cyclin B3 siRNA *(B)* or synchronized in mitosis with nocodazole *(C)*. α-Tubulin serves as a loading control. **(D)** *(top)* Schematic illustrating the strategy utilized for cell synchronization and collection of extracts. *(bottom)* Immunoblots of synchronized HeLa cell extracts. α-Tubulin serves as a loading control. **(E)** Immunoblot of 3xFlag-cyclin B3 complexes immunoprecipitated from mitotic HeLa cells, showing interaction of cyclin B3 with CDK1 and CDK2. **(F)** *(top)* Schematic illustrating strategy used for imaging HeLa FUCCI cells. *(bottom)* Time-lapse sequences of HeLa cells labelled with the FUCCI marker, undergoing cell cycle progression. G1 cells are labelled in magenta, S and G2 cells are in green. Cells were also followed by DIC microscopy (grey). Scale bar, 10 µm. **(G)** Quantification of cell cycle stage durations from HeLa FUCCI movies. **(H)** Time-lapse sequences of HeLa cells expressing GFP-tagged PCNA and mRFP-tagged Histone H2B, undergoing cell cycle progression. The disappearance of PCNA foci marks the beginning of G2 and nuclear envelope breakdown (NEBD) marks the beginning of mitosis. Scale bar, 10 µm. **(I)** Quantification of cell cycle stage durations from HeLa PCNA movies. **** represents *P < 0.0001* from Mann-Whitney tests; non-significant (n.s.) is *P > 0.05*.

### Cyclin B3 is not required for interphase cell cycle progression

Having established that cyclin B3 is expressed in mitotic cells, we next evaluated its functionality. Using the FUCCI reporter system (Koh et al., 2017), we found that cyclin B3 depletion through siRNA had no significant effect on G1 duration, nor on the combined duration of the S and G2 phases (**Fig. 1F&G**). Moreover, using HeLa cells stably expressing GFP-tagged PCNA to measure G2 progression (Leonhardt et al., 2000) (**Fig. 1H**), we found that cyclin B3 depletion had no effect on G2 duration (**Fig. 1I**). Similar observations were derived using the osteosarcoma cell line U2-OS (**Fig. S1C**). Collectively, these results indicate that, cyclin B3 is not required for mitotic entry or for interphase cell cycle progression.

### Cyclin B3 ensures timely mitotic progression

Interestingly, despite not being required for interphase, cyclin B3 depletion significantly impaired cells’ ability to progress through mitosis (**Fig. 1G&I**). To quantify this, we filmed HeLa cells stably expressing mRFP-tagged histone H2B (**Fig. 2A**). While control cells completed mitosis in ∼55 minutes, cyclin B3 depleted cells extended this timing to ∼200 minutes (**Fig. 2B**). This extension in mitotic duration was suppressed by expression of an siRNA-resistant wild-type cyclin B3, but not a CDK binding mutant (CB^mut^: Y1129, I1133, Y1136) (Goda et al., 2001; Lara-Gonzalez et al., 2024) (**Fig. 2C&D; S1D&E**). To determine the specificity of cyclin B3 in promoting mitotic progression, we generated HeLa cells expressing doxycycline-inducible cyclin B1 (**Fig. 2E**). We found that the effect of cyclin B3 depletion could not be suppressed by cyclin B1 overexpression (**Fig. 2F**), establishing that the observed effects are specific to cyclin B3. We conclude that cyclin B3 promotes efficient mitotic progression through its ability to bind and activate CDK1/2.

**Figure 2.**
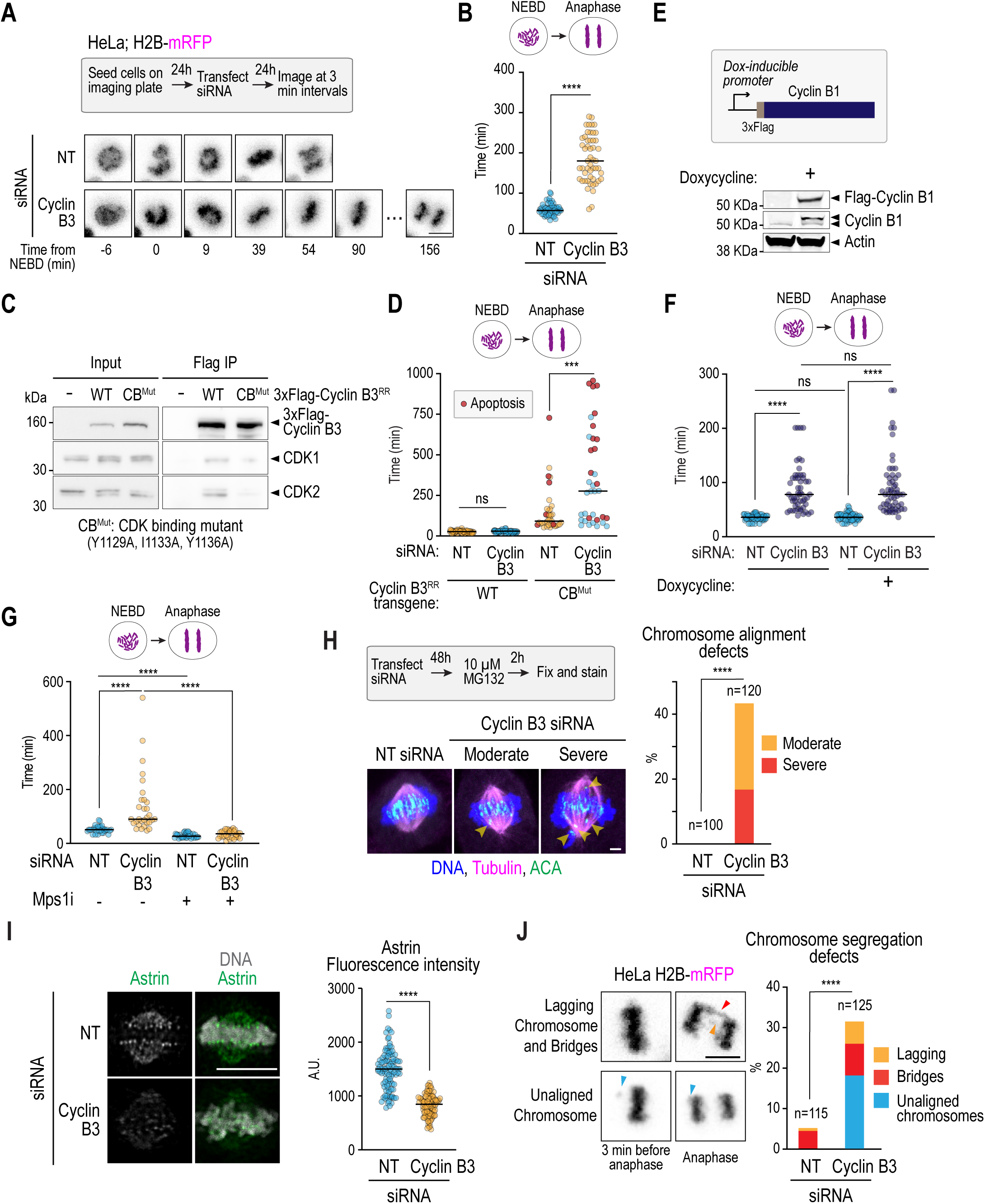
Cyclin B3-CDK1/2 ensures timely mitotic progression and fidelity. **(A)** Time-lapse sequence of HeLa cells expressing mRFP-tagged histone H2B and treated with either non-targeting siRNA (NT) or cyclin B3 siRNA. Numbers at the bottom indicate the time elapsed from nuclear envelope breakdown (NEBD) in minutes. **(B)** Quantification of the interval from NEBD to Anaphase onset under the indicated conditions. **(C)** Immunoblot of 3xFlag-cyclin B3 immune complexes isolated from mitotic HeLa cells, using either cyclin B3 wild-type (WT) or a CDK binding mutant (CB^Mut^) and probed for CKD1 and CDK2. **(D)** Quantification of the interval from NEBD to Anaphase onset for the indicated conditions. Red dots indicate cells that underwent apoptosis at the end of the experiment. Note that cells depleted of cyclin B3 and expressing cyclin B3 CB^Mut^ prolonged mitosis to a greater extent than depletion of cyclin B3 alone, suggesting that CB^Mut^ cyclin B3 may act in a dominant-negative manner. **(E)** *(top)* Schematic illustrating the construct used for the doxycycline-induced overexpression of 3xFlag-tagged cyclin B1. *(bottom)* Immunoblot showing 3xFlag-cyclin B1 overexpression. α-Tubulin serves as a loading control. **(F)** Quantification of the interval from NEBD to Anaphase onset for the indicated conditions. **(G)** Quantification of the interval from NEBD to Anaphase onset under the indicated conditions. Mps1i refers to NMS-P715. **(H)** *(left)* Immunofluorescence images of HeLa cells depleted of cyclin B3 and stained with DAPI (blue), α-Tubulin (magenta) or centromeres (ACA, green) and illustrating examples of chromosome alignment errors. Scale bar, 2 µm. *(right)* Quantification of chromosome alignment errors for the indicated conditions. **(I)** *(left)* Immunofluorescence images of HeLa cells depleted of cyclin B3 and stained with DAPI (blue) and Astrin (green). *(right)* Quantification of Astrin kinetochore fluorescence levels. For each condition, 10 kinetochores were scored in 10 individual cells, for a total of 100 kinetochores per condition. **(J)** *(left)* Representative images of HeLa cells expressing mRFP-tagged histone H2B and imaged through time-lapse microscopy. Examples of unaligned chromosomes and lagging chromosomes detected are shown. Scale bar, 10 µm. *(right)* Quantification of mitotic chromosome segregation errors under the specified conditions. *n* represents the number of cells analyzed. Graphs from panels *B, D, F, G* and *H* were analyzed using Mann-Whitney tests and panels *I* and *J* were analyzed using Chi square analysis. **** represents *P < 0.0001*; *** represents *P < 0.001*; non-significant (n.s.) is *P > 0.05*.

### Cyclin B3 is required for the formation of proper kinetochore-microtubule attachments in mitosis

Delays in mitotic duration may arise from activation of the spindle assembly checkpoint (SAC), a surveillance mechanism that delays mitotic progression when kinetochores are devoid of microtubule attachments (Hayward et al., 2019; Lara-Gonzalez et al., 2021; McAinsh and Kops, 2023). To test this, we treated cyclin B3 depleted cells with a small molecule inhibitor of the SAC kinase Mps1, which abolished the cyclin B3 depletion – induced mitotic delay (**Fig. 2G**). These findings suggest that cyclin B3 is required for normal kinetochore-microtubule attachments in mitosis. In agreement with this, cyclin B3 depletion resulted in ∼45% of cells exhibiting moderate (one unaligned chromosome) to severe (two or more unaligned chromosomes and/or polar chromosomes) alignment defects (**Fig. 2H)** and resulted in a substantially broader metaphase plate (**Fig. S2A**). Moreover, cyclin B3-depleted cells showed reduced kinetochore levels of Astrin, a component of the Astrin-SKAP complex that marks mature kinetochore-microtubule attachments (Ying et al., 2020) (**Fig. 2I**). Live imaging also revealed that ∼18% of cyclin B3 depleted cells entered anaphase with unaligned chromosomes and ∼16% exhibited lagging chromosomes or chromosome bridges (**Fig. 2J; S2B**). These findings reveal that cyclin B3 plays a critical role in ensuring proper kinetochore-microtubule attachments in mitosis and that its loss delays mitosis by activating the SAC. In addition, attachment errors resulting from loss of cyclin B3 lead to increased chromosome segregation errors during anaphase. Consistent with cyclin B3 being required for mitotic fidelity, we found that cyclin B3 depletion led to reduced colony formation and growth rates (**Fig. S2C&D**).

### Cyclin B3-CDK1/2 phosphorylates a wide range of mitotic substrates

Having established that cyclin B3-CDK1/2 contributes to mitotic timing and fidelity, we next sought to identify its substrates using mass spectrometry. To the best of our knowledge, the only confirmed substrate of cyclin B3 corresponds to Emi2/XErp1 (Bouftas et al., 2022), which alone may not fully account for the mitotic defects observed upon cyclin B3 depletion. Using an inducible cyclin B3 CRISPR-Cas9 system (**Fig. S3A-C**), we generated mitotic extracts from control and cyclin B3 depleted cells (**Fig. 3A**), enriched for phospho-peptides and analyzed through shotgun proteomics. Our analysis identified 1528 phopho-sites, matching 1203 unique proteins, significantly reduced abundance upon cyclin B3 loss (**Fig. 3B, S3C & D; Table S1**). PhosphoSitePlus analysis indicated that 85 of these proteins are predicted CDK1 and 65 are CDK2 targets (**Fig. 3C**). In accordance, ∼41% of identified phospho-peptides contained a consensus CDK phosphorylation site ([S/T]-P) (**Fig. 3B**), which is suggestive of direct targets of the cyclin B3-CDK1/2 complex in mitosis. Gene Ontology (GO) analysis revealed significant enrichment for cell cycle regulators, as well as proteins involved in processes such as chromatin remodeling, chromatin organization and RNA metabolism (**Fig. 3D**). To identify mitotic processes regulated by the cyclin B3-CDK1/2 complex, we performed clustering analyses on mitotic regulators (**Fig. 3E**). This revealed a large cluster of proteins involved in sister chromatid segregation, including kinetochore proteins such as BUB1, KNL1, CENPF, and NUF2. Smaller clusters were associated with other chromosomal functions, such as nucleotide excision repair, DNA damage response, and activation of pre-replicative complexes. Additionally, spindle-associated factors such as TPX2, PRC1 KIF20A were identified (**Fig. 3E**). A top candidate that emerged from our analysis corresponded to CEP192 (**Table S1**), a pericentriolar matrix (PCM) component that promotes mitotic spindle assembly (Varadarajan and Rusan, 2018). Interestingly, CEP192 levels were reduced at centrosomes following cyclin B3 depletion (**Fig. 3F**), suggesting that CEP192 phosphorylation by cyclin B3-CDK1/2 ensures its PCM recruitment. Overall, our analysis suggests that cyclin B3, in complex with CDK1/2, phosphorylates a wide range of mitotic substrates to ensure mitotic fidelity and mitotic progression.

**Figure 3.**
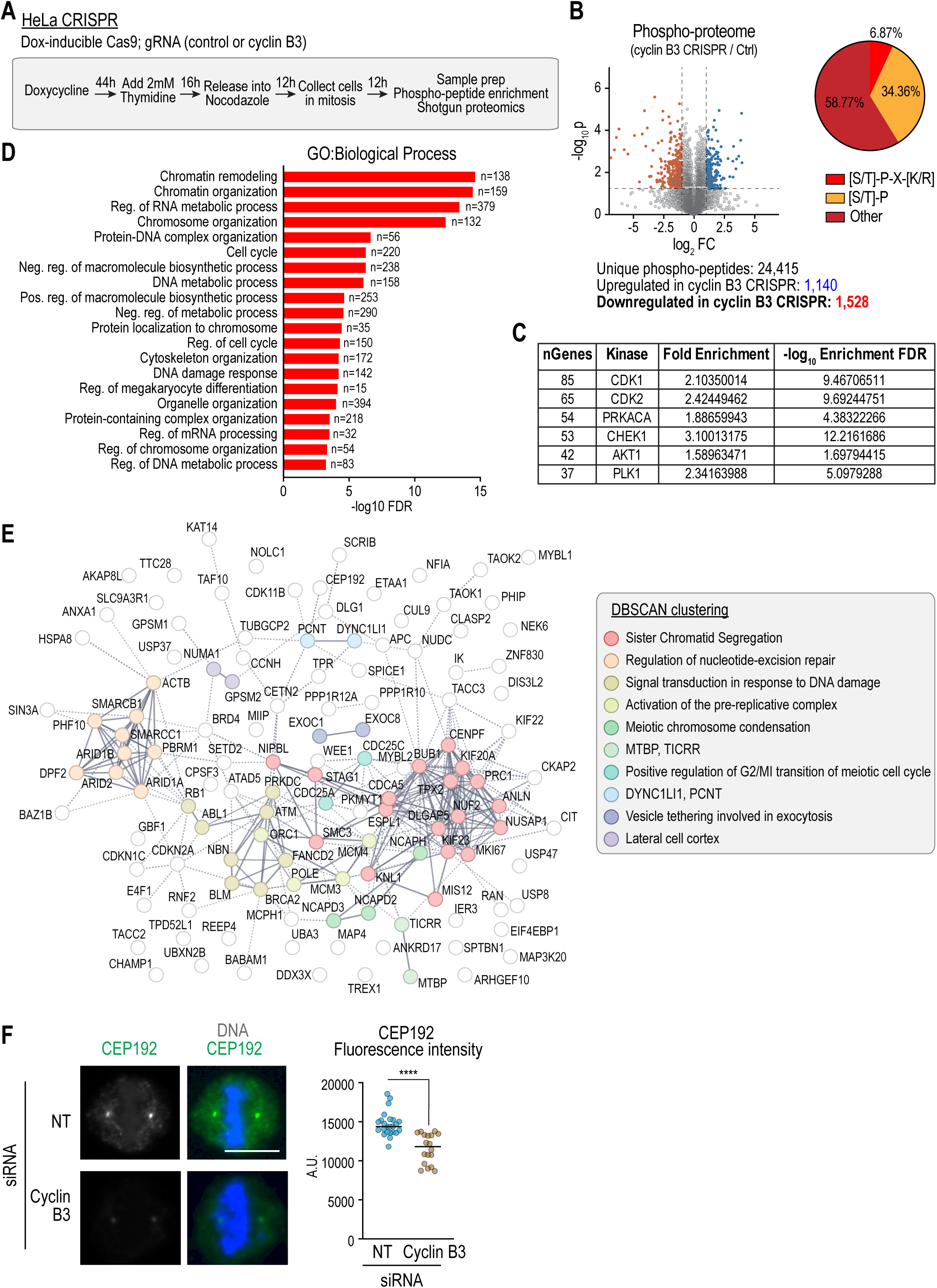
Identification of cyclin B3 mitotic substrates through mass spectrometry. **(A)** Strategy utilized for the generation of mitotic extracts depleted of cyclin B3 for the identification of differentially phosphorylated peptides through phospho-proteomic analysis. See *Fig. S3A&B* for the generation and characterization of doxycycline-induced CRISPR-Cas9 lines **(B)** *(left)* Volcano plot illustrating differentially phosphorylated phospho-peptides after normalization by protein abundance. *(right)* Pie chart illustrating the phospho-peptide signatures for the putative cyclin B3-CDK1/2 substrates identified through mass spectrometry. **(C)** PhosSitePlus analysis of potential cyclin B3-CDK1/2 substrates showing enrichment in CDK1 and CDK2 substrates. **(D)** GO Term analysis of potential cyclin B3-CDK1/2 substrates identified in our study, showing enrichment of proteins with chromosome segregation and cell cycle regulation functions. Proteins were ranked by –log_10_FDR (false discovery rate). **(E)** STRING analysis and DBSCAN clustering of proteins marked as “mitotic cell cycle” from our GO Term analysis, showing clusters of proteins with common functions. **(F)** *(left)* Immunofluorescence images of HeLa cells depleted of cyclin B3 and stained with DAPI (blue) and CEP192 (green). *(right)* Quantification of CEP192 centrosome fluorescence levels. Scale bar, 10 µm. **** represents *P < 0.0001* from Mann-Whitney tests; non-significant (n.s.) is *P > 0.05*.

### Cyclin B3 overexpression accelerates mitotic progression

Having established that cyclin B3 depletion extends mitotic duration in cancer cells, we tested the effects of its overexpression. While cyclin B3 overexpression had no significant effect on interphase cell cycle duration (**Fig. S4C**), total mitotic duration was reduced from ∼55 to ∼28 minutes (**Fig. 4A&B**) largely due to a reduction in the timing between NEBD and metaphase (**Fig. S4A**), suggesting increased chromosome alignment efficiency. Consistent with this, the metaphase plate width was significantly narrower in cyclin B3-overexpressing cells relative to controls (**Fig. S4B**). Moreover, overexpression of wild-type cyclin B3, but not the CDK binding mutant, resulted in accelerated mitosis (**Fig. 4B**), confirming that this effect is mediated by cyclin B3’s ability to activate CDK1/2. Collectively, these findings indicate that cyclin B3 enhances the fidelity and efficiency of mitotic progression.

**Figure 4.**
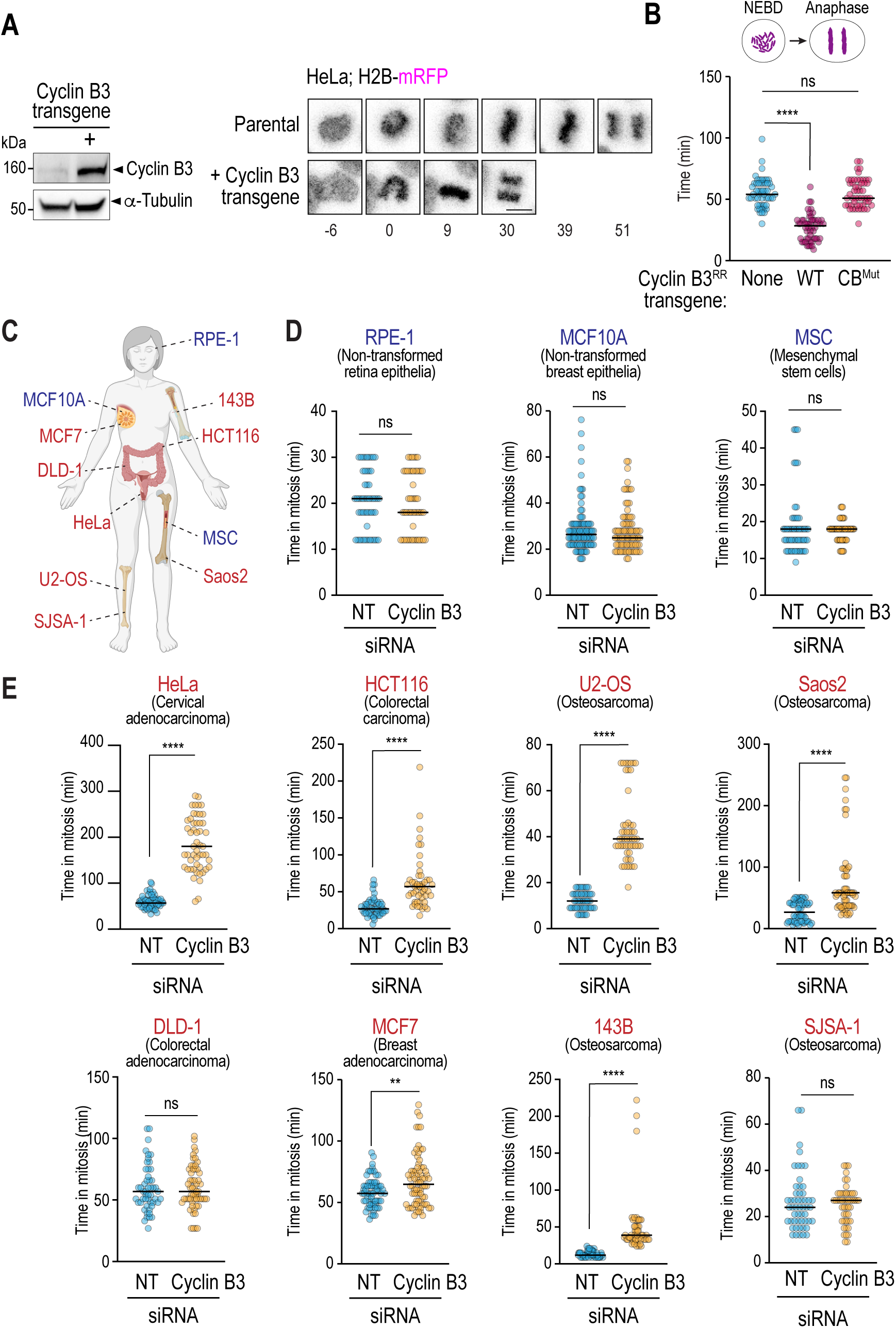
Cyclin B3 accelerates mitotic progression in cancer cells. **(A)** *(left)* Immunoblot showing cyclin B3 overexpression. α-Tubulin serves as a loading control. *(right)* Time-lapse sequence of HeLa cells expressing mRFP-tagged histone H2B, either parental or cyclin B3 overexpressing. Numbers at the bottom indicate the timing from nuclear envelope breakdown (NEBD) in minutes. Scale bar, 10 µm. **(B)** Quantification of the interval from NEBD to Anaphase onset for the indicated conditions. **(C)** Cartoon illustrating the tissue origin of the cell lines used in this study (created with BioRender). **(D)**&**(E)** Quantification of mitotic duration for the specified non-transformed *(B)* or cancerous *(C)* cell lines. For RPE-1, HeLa, HCT116, U2-OS and DLD-1, mitotic duration corresponds to the timing from NEBD to Anaphase onset and was determined using cells expressing mRFP-tagged histone H2B. For MCF10A, MSC, Saos2, MCF7, 143B and SJSA-1, mitotic duration corresponds to the timing from cell rounding to anaphase onset and was determined using DIC imaging. The HeLa mitotic timing data presented in this figure is the same as Fig. 2B. **** represents *P < 0.0001* from Mann-Whitney tests; ** represents *P < 0.01*; non-significant (n.s.) is *P > 0.05*.

### Cyclin B3 is crucial for normal mitotic duration in cancer cells

Overall, our data established that cyclin B3 is required for mitotic progression and fidelity in both HeLa and U2-OS cells, likely due to its role as an activator of CDK1/2. This finding was, however, seemingly at odds with the notion that vertebrate cyclin B3 does not appear to play a significant role in cell cycle control outside of female meiosis (Fatemi et al., 2021; Karasu and Keeney, 2019; Rezaei et al., 2022; Wang et al., 2023). To address this discrepancy, we expanded our analysis of cyclin B3 mitotic phenotypes across a panel of cancer and non-cancerous cell lines (**Fig. 4C; Table S2**). Strikingly, depletion of cyclin B3 in non-transformed lines did not cause any significant extension of mitotic duration, while the majority of cancer cell lines recapitulated the prolonged mitotic timing (**Fig. 4D&E; S5A**). Moreover, while cyclin B3 mRNA was detected in all cell lines (**Fig. S5A**), protein levels were higher in cancer cells (**Fig. S5B**). Collectively, these data suggest that expression of cyclin B3 in cancer ensures their normal progression through mitosis.

### A role for a meiotic B-type cyclin in driving mitotic progression and fidelity in cancer

The work presented here uncovers an unexpected role for cyclin B3, a meiotic B-type cyclin, as a key driver of mitotic progression and fidelity in cancer cells by ensuring efficient kinetochore-microtubule attachments (**Fig. 5**). Failure of cyclin B1 to compensate for the loss of cyclin B3 and its inability to accelerate mitosis suggest that cyclin B3 has specific functions that cannot be compensated by other cyclins. Whether this specificity arises from cyclin B3 recognizing specific mitotic substrates (Ord et al., 2025), serving as a more potent CDK activator (Lara-Gonzalez et al., 2024) or targeting a broader number of substrates remains to be determined.

**Figure 5.**
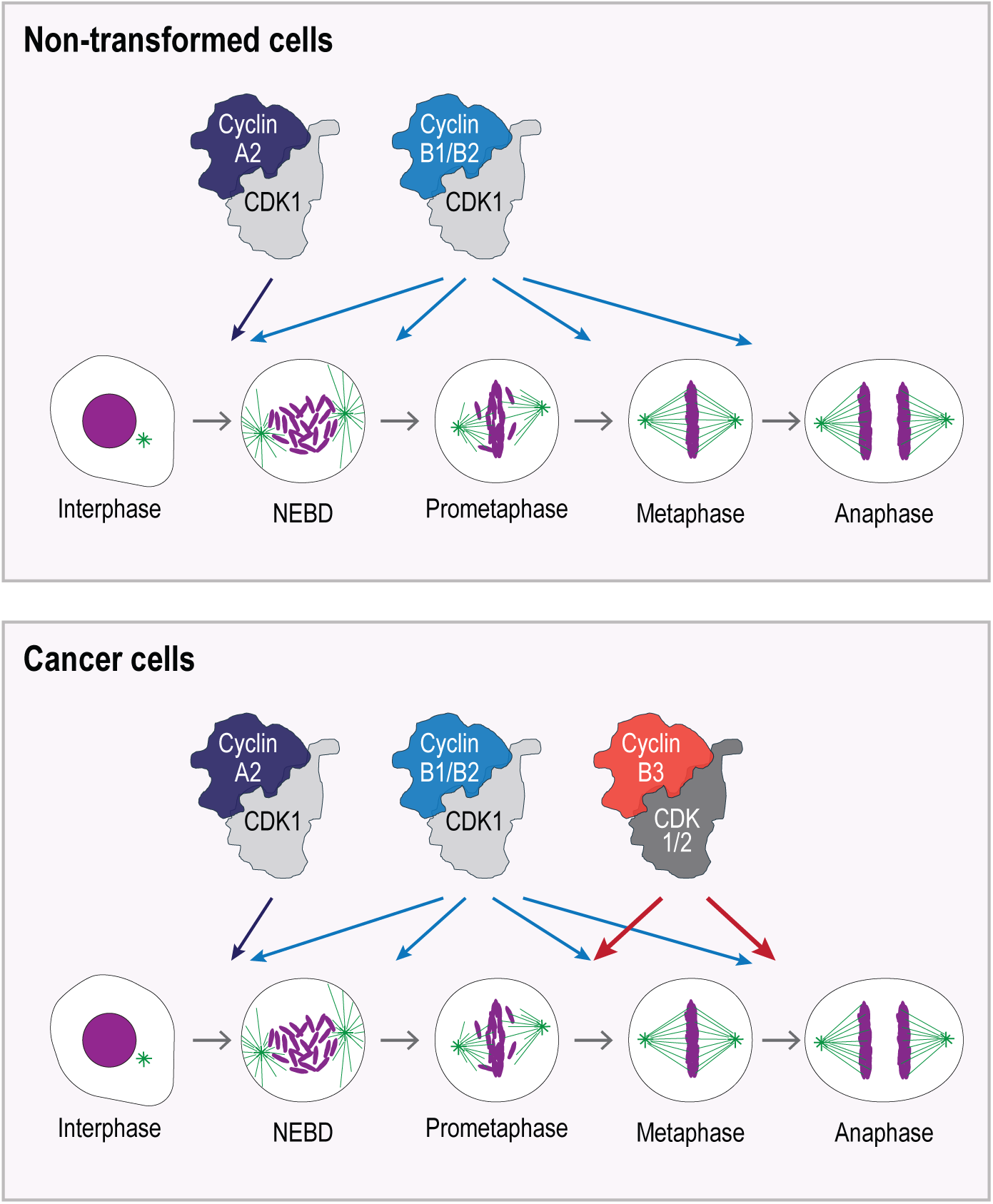
Cyclin B3-CDK1/2 ensures mitotic progression in cancer cells. In non-transformed somatic cells, canonical B-type cyclins (cyclins B1 and B2) drive mitotic progression. In addition, the cyclin A2-CDK1 complex promotes the transition from interphase into mitosis, either by cooperating with cyclins B1 and B2 or by activating the Greatwall-ENSA pathway (Crncec et al., 2025; Hegarat et al., 2020). In cancer cells, cyclin B3 forms active complexes with CDK1 and CDK2 to promote faithful kinetochore-microtubule attachments and accurate chromosome segregation, adding an additional level of cell cycle control. We propose that cyclin B3-CDK1/2 protects cancer cells against severe mitotic errors that may push them towards cell death.

Our findings are consistent with Cancer Dependency Map (DepMap) data identifying cyclin B3 as an essential gene in ∼20% of cell lines through shRNA-mediated depletion (Shimada et al., 2021; Tsherniak et al., 2017). While no cancer type emerged as particularly sensitive from our analyses, we note that Ewin-like sarcomas frequently overexpress BCOR-cyclin B3 fusions that are thought to provide them with a proliferative advantage (Argani et al., 2017; Cidre-Aranaz et al., 2022; Pierron et al., 2012). Our finding that cyclin B3 overexpression improves mitotic quality in cancer suggests that cellular transformation or associated alterations in cell cycle regulation may increase their dependency on cyclin B3 for efficient mitotic progression. Targeting meiotic pathways in cancer, including cyclin B3-CDK1/2, could provide a foundation for the development of novel anti-cancer therapies with reduced toxicities.

## MATERIALS AND METHODS

### Cell lines and cell culture

All cell lines were cultured in humidified incubators at 37**°**C with 5% CO2.

HeLa (CRM-CCL-2), MCF7 (HTB-22) and HEK 293T (CRL-3216) cells were grown in Dulbecco’s Modified Eagle Medium (DMEM) (Gibco) supplemented with 10% Fetal Bovine Serum (FBS), 100 units/mL penicillin, 100 μg/mL streptomycin, 4.5 g/L D-Glucose, 4 mM L-glutamine, 1 mM sodium pyruvate, and GlutaMAX. HCT-116 (CCL-247), Saos-2 (HTB-85), U2-OS (HTB-96) cells were grown in McCoy’s 5A Medium (Gibco) supplemented with 10% Fetal Bovine Serum (FBS), 100 units/mL penicillin, 100 μg/mL streptomycin, and 1.5 mM L-glutamine. RPE-1 (CRL-4000) cells were grown in DMEM-F12 medium supplemented with 10% Fetal Bovine Serum (FBS), 100 units/mL penicillin and 100 μg/mL streptomycin. SJSA-1(CRL-2098) and DLD-1 (CCL-221) were grown in RPMI-1640 medium (Gibco) supplemented with 2 mM L-glutamine, 10 mM HEPES, 1 mM sodium pyruvate, 4500 mg/L glucose, and 1500 mg/L sodium bicarbonate, 10% Fetal Bovine Serum (FBS), 100 units/mL penicillin and 100 μg/mL streptomycin. Mesenchymal Stem Cells (MSCs; (Mihara et al., 2003)) were grown in DMEM supplemented with 10% Fetal Bovine Serum (FBS), 100 units/mL penicillin, 100 μg/mL streptomycin. 143B (CRL-8303) were grown in MEM supplemented with 0.015 mg/mL 5-bromo-2’-deoxyuridine, 10% Fetal Bovine Serum (FBS), 100 units/mL penicillin and 100 μg/mL streptomycin. MCF10A (CRL-10317) was grown in MEBM® Basal Medium (Lonza #CC-3151) with MEGM^®^ SingleQuots^®^ Supplements (Lonza #CC-4136) and Cholera Toxin (Sigma #NC1153072).

All cell lines were authenticated through short tandem repeat (STR) DNA profiling (UC Berkeley DNA Sequencing Facility) and confirmed negative for mycoplasma contamination using the PlasmoTest kit (Invivogen #rep-pt1).

### Antibody production

To generate an anti-cyclin B3 antibody, a cDNA fragment encoding for cyclin B3 amino acids 1-80 was expressed in *E. coli* as a fusion to maltose-binding protein (MBP) and the resulting purified protein was used to immunize rabbits (Labcorp). Antibodies were affinity purified from the serum of immunized animals using a fragment containing amino acids 1–80 of cyclin B3 fused to GST (Desai et al., 2003). Antibody specificity was tested through immunoblotting, using HeLa cell lysates depleted of by cyclin B3 through siRNA (**Fig. 1B**).

### siRNA Transfection

For siRNA transfections, cells were seeded onto 6-well plates with appropriate media without antibiotics and transfected with either a non-targeting siRNA (Horizon Discovery #D-001810-10-20) or a cyclin B3 SMARTpool siRNA (Horizon Discovery #L-003208-00-0005 (GUCUCAAGGCUGUGUAUUA, GCGCAGAUAUGCUAGGUGU, UGAACAAACUGCUGACUUU, CAACUCACCUCGUGUGGAU)) using Lipofectamine RNAiMAX (Thermo Fisher Scientific #13778-075), according to the manufacturer’s protocol. Knockdown efficiency was assessed by quantitative PCR 48h after transfection.

### Cell cycle synchronization

For synchronizing cells in mitosis, cells were treated with 1 µM Nocodazole for 16 hr. For the experiments in **Fig. 1C**, cells were treated with 2 mM Thymidine (Sigma # T9250-1G) for 16 hr and then released into fresh media for 9 hr after washing thrice with PBS. Cells were then treated again with 2 mM Thymidine for 16h and then released into media containing 1 µM Nocodazole after washing thrice with PBS.

### Crystal violet staining

Cells were fixed for 10 minutes in 100% methanol for 10 minutes at room temperature and stained with 1% Crystal violet for 30 minutes. The cells were then washed with PBS 3 times, dried and imaged.

### Preparation of cell lysates for immunoblotting

Cells were seeded onto 6 well plates approximately 24h prior to treatment. After treatment, media in the wells was removed and cells were lysed directly in 300 µL of 2X Sample buffer (116.7 mM Tris-HCl pH 6.8, 3.3% SDS, 200 mM DTT, 10% glycerol, bromophenol blue), followed by heating at 100**°**C for 10 minutes.

### Immunoprecipitation

Cells were resuspended in 400 µL of lysis buffer (10 mM Tris-HCl pH 7.5, 100 mM NaCl, 0.1% Triton X-100, 1 mM EGTA, 1 mM EDTA, 1 mM DTT, 20 mM β-glycerol phosphate, 10 mM sodium fluoride, 0.2 mM PMSF and EDTA-free protease inhibitor cocktail (Sigma #11697498001)). Cell suspensions were rotated for 30 minutes at 4°C before centrifugation at 13,000 rpm for 30 min at 4°C. For input, 25 μL of supernatant was taken and mixed with 25 μL of 4X SDS sample buffer and boiled for 10 minutes. The remaining supernatant was incubated with anti-FLAG magnetic beads (Millipore-Sigma # M8823-1ML), overnight at 4°C on an end-over-end rotator. After incubation, beads were washed 3 times with lysis buffer, lysed in 50 μL of 2X SDS-sample buffer and boiled for 10 minutes.

### Expression and immunoprecipitation of proteins from HEK293T cells

cDNAs encoding for cyclin B3, cyclin B1 and CDK-1 were cloned onto pcDNA3.1-based vectors encoding for 5xMyc or 3xFlag tags, respectively (**Table S3**). For expression, HEK293T cells were grown in10 cm dishes per condition and transfected at 60% confluency with 10 µg of DNA constructs per plate using Lipofectamine 3000 Transfection Reagent and serum-free DMEM medium, according to product guidelines (Thermo Fisher Scientific #L3000001). Transfected cells were incubated for 48 h at 37°C and 5% CO2. Lysis and immunoprecipitation was performed as described above, using Pierce™ Anti-c-Myc Magnetic Beads (Thermo Fisher Scientific #88842).

### Immunoblotting

For immunoblotting, samples were loaded onto 4-20% Bis-Tris Gels (GenScript, #M00657). Proteins were transferred to nitrocellulose membranes using the Trans-Blot Turbo Transfer System (Bio-Rad, #1704158). Subsequently, membranes were blocked with intercept blocking buffer (LicorBio #927-60001) and incubated overnight with primary antibody diluted in intercept blocking buffer. Membranes were then washed with 0.1% TBS-T (20 mM Tris-HCl, 150 mM NaCl, 0.1% Tween 20) before a one-hour incubation with secondary antibody in intercept blocking buffer, at room temperature. Membranes were washed with TBS-T and developed in either a LI-COR imaging system for fluorescent secondary antibodies or a BioRad ChemiDoc for HRP conjugated secondary antibodies.

Antibodies used for immunoblotting were: rabbit anti-Cyclin B3 (this study), mouse anti-phospho-Histone H3 (Ser10) (clone CMA312, Millipore #05-1336), rabbit anti-Cyclin B1 (Cell Signaling Technology #4138), mouse anti-Cdc2 p34 (Santa Cruz #sc-54), mouse anti-Cdk2 (Clone D-12, Santa Cruz #sc-6248), mouse anti-Flag (clone M2, Sigma-Aldrich #F3165), mouse anti-α-tubulin (clone DM1A, Millipore #T6199), mouse anti-Actin (clone C4, Millipore #MAB1501), 800CW goat anti-rabbit IgG (LI-COR #926-32211), Peroxidase AffiniPure Goat Anti-Rabbit IgG (Jackson Immunoresearch #111-035-003) Peroxidase AffiniPure® Goat Anti-Mouse IgG, light chain specific (Jackson Immunoresearch #115-035-174) and Alexa Fluor 680 goat anti-mouse IgG light chain specific (Jackson Immunoresearch #115-625-174).

### Generation of transgenic cell lines

Transgenic cell lines were generated by lentiviral transduction. Plasmids used for lentivirus generation are listed in **Table S3**.

Lentiviral particles were generated using the Lenti-X packaging single shots with VSV-G (Takara Bio #631278) in Lenti-X 293T cells in DMEM media, according to the manufacturer’s instructions. Media containing viruses was collected 48h after transfection, spun down at 1000 g for 10 minutes to remove cell debris. 60-80% confluent cells seeded onto a 10 cm dish were then transduced using 3 mL of virus containing media and 5μg/mL Polybrene. After 48h, parental and viral infected cells were treated with the appropriate antibiotic – 5 μg/mL Blasticidin S (Thermo Fisher Scientific #AAJ67216XF) or 5 μg/mL Puromycin (Thermo Fisher Scientific #J67236XF) – to select for stably integrated cells. Antibiotic selection was terminated once the treatment eliminated all cells in a non-transduced control plate that was tested simultaneously. The surviving infected cells were pooled and cultured. To generate lines expressing H2B-mRFP, cells were sorted using a FACS ARIA II sorter (UC Irvine Stem Cell Research Centre Flow & Mass Cytometry Core), selecting for cells with intermediate mRFP fluorescence levels.

### Generation of inducible CCNB3 CRISPR-Cas9 cell lines

Hela cells were engineered to stably express Tet inducible Cas9 and gRNA targeting Cyclin B3 by lentiviral transduction of the TLCV2 backbone (Addgene plasmid #87360) with or without Cyclin B3 gRNA (gRNA1 sequence: AGTGGTAGGCTTCTTAATGG gRNA2 sequence: TTAAGCATACCAACAAAAGT). Expression of Cas9 was induced by treatment with 0.5μg/mL Doxycycline Hyclate (Dox) (Sigma Cat #D5207-1G) for 48 hours, followed by washing out for 24 hours prior to analysis. Unless otherwise indicated, Dox was washed off 48h after addition.

### Live imaging microscopy

For live imaging microscopy, cells were seeded into 18 chamber glass bottom slides (Ibidi #81817). Live imaging was done overnight in an environmentally controlled chamber at 37°C and 5% CO_2_. For experiments in **Fig. 2G**, cells were treated with DMSO or 200 nM NMS-P715 just before imaging. Imaging was performed using Zeiss LSM 780 or Zeiss LSM 980 confocal microscopes with 488 and 561 nm laser lines using the ZEN black (LSM 780) or ZEN blue (LSM 980) software. Transmitted light images were also collected. Standard PMT detectors were used on LSM 780 and GaAsP (Gallium Arsenide Phosphide photocathode) PMTs were used on LSM 980. Images were acquired with a Zeiss 20x objective and were captured every 3 or 10 min in 3 x 2.5 μm z-sections.

### Immunofluorescence

For immunofluorescence, cells growing on glass coverslips were fixed with 4% or 1% *(Fig. 2I)* formaldehyde (in PBS) for 30 minutes at room temperature. Cells were then washed 3 times 10 minutes each with PBS-T (0.3% Triton-X 100 in PBS), blocked in PBS-T with 3% BSA and incubated overnight at 4°C in primary antibody diluted in blocking buffer. After washing 3 times with PBS-T, cells were incubated for 30 minutes with fluorescently labelled secondary antibodies. Coverslips were mounted in ProLong Diamond with DAPI (Thermo Fisher Scientific #P36966) and imaged on a Zeiss LSM 780 or Zeiss LSM 980 microscope using a 63X objective. For fluorescence intensity quantifications in *Fig. 2I* and *3F*, images were captured on a Nikon spinning disk microscope system coupled with a CSU-X1 spinning disk unit (Yokogawa) and using a 60X objective.

Antibodies used for immunofluorescence were: anti-centromere antibody (ACA) (Antibodies Incorporated #SKU: 15-235), anti-Cep192 (Sigma-Aldrich #HPA039392) anti-α-tubulin (clone DM1A, Millipore #T6199), anti-Astrin (Bio-Techne #NB100-74638), anti-Flag (clone M2, Sigma-Aldrich #F3165), Alexa Fluor 488 AffiniPure Goat Anti-Human IgG (H+L) (Jackson Immunoresearch #109-545-003), Cy™5 AffiniPure® Donkey Anti-Mouse IgG (H+L) (Jackson Immunoresearch #715-175-151) and Cy™2 AffiniPure™ Donkey Anti-Rabbit IgG (H+L) (Jackson Immunoresearch #711-225-152).

### Imaging analysis

Imaging analysis was performed on ImageJ, using maximum intensity projections. Mitotic timing was determined using H2B-mRFP (RPE1, HeLa, U2OS, HCT116, DLD-1) or transmitted light (MCF 10A, MSC, Saos2, MCF7, 143-B, SJSA1). For cells expressing H2B-mRFP, nuclear envelope breakdown (NEBD) was defined as the point where diffused H2B-mRFP signal can be seen to condense, and chromosomes can clearly be seen to move. Metaphase was defined as the point at which the aligned chromosome width (metaphase plate) was the smallest. Anaphase was defined as the point where chromosomes can clearly be seen to separate. Mitotic time from transmitted light movies were defined as the time from cell rounding to anaphase.

For cell cycle timing using FUCCI, time in G1 was defined as the time between start of mKO2 expression and the onset of mAG expression. The brief time where both fluorophores are expressed together was considered to be S/G2. Timing of S/G2 was defined as the time between the start of mAG expression and cell rounding. Time in mitosis was defined as time between cell rounding and anaphase as observed from transmitted light images. For G2 timing analysis from GFP-PCNA, G2 onset was defined as the first time point after the disappearance of PCNA foci. Time in G2 was defined as the time between disappearance of PCNA foci and NEBD, as observed with H2B-mRFP.

Metaphase plate width was measured from movies using H2B-mRFP. Using FIJI/Image J, a bounding box was drawn to include all mRFP signal 2 time points/frames (6 minutes) before anaphase. The width of the bounding box was measured as metaphase plate width. Chromosome alignment defect was considered moderate if the ACA signal of one chromosome was misaligned but still close to the metaphase plate (as indicated in *Fig. 3B,* moderate panel). Chromosome alignment defects were considered severe if ACA signal was distributed throughout the spindle, at spindle poles or separate / unattached to the spindle (as indicated in *Fig. 3B*, severe panel).

To quantify astrin signal intensities at kinetochores, we used the two-box method (Hoffman et al., 2001). Using FIJI/ImageJ, the first box was drawn around the kinetochores, and the integrated intensity was measured. This box was then expanded by 2 pixels in each direction, and the average intensity in the region between the expanded box and the first box was used to measure the local background. This was then subtracted from the integrated density of the first box (drawn around the kinetochores) and the resulting intensity was plotted in *Fig. 2I*.

### RNA extraction, cDNA synthesis and Quantitative PCR

RNA was extracted from cells using TRIzol (Thermo Fisher Scientific #15596018) according to manufacturer’s instructions. cDNA was synthesized using qScript Ultra SuperMix (Quantabio # 95217-025) and quantitative PCR (qPCR) was performed using PerfeCTa SYBR Green FastMix Low ROX (Quantabio # 95074-250). The following primers were used for qPCR: β-actin: Fw-CCCGCCGCCAGCTCA, Rev-ACGATGGAGGGGAAGACGG; Cyclin B3: Fw-CAAGGGAACACCAAAGGAGAT, Rev-TCTGTAAGTATAAACTGTTCCTCTC. Relative mRNA levels were quantified using the formula RE = 2^-ΔΔ^Ct method.

### Protein Extraction and Sample Preparation for Mass Spectrometry

HeLa cell pellets were lysed in 1 mL of 50 mM ammonium bicarbonate, pH 8.5 buffer containing 4 M urea, 0.5% SDS, 1 mM DTT, 100 mM NaCl, and supplemented with LPA (leupeptin, pepstatin, aprotinin) protease and phosphatase inhibitors. Samples were first needle-lysed by 10x passages through a 22-gauge syringe needle on ice, then further disrupted by sonication using a Diagenode BioRuptor® Pico (5×30s). The resulting lysates were then centrifuged at 15,000 rcf for 15 min at 4°C to pellet insoluble material and the clarified supernatants were then measured by BCA protein assay.

### Filter-Aided Sample Preparation Reduction, Alkylation, and Tryptic Digestion

Protein digestion was performed using a filter-aided sample preparation (FASP) approach. For each sample, clarified lysate was diluted to reduce SDS concentration to 0.1% and then transferred to and distributed across four Microcon® 30-kDa centrifugal filters (NMWCO 30 kDa, Ultracel® regenerated cellulose membrane, Sigma-Aldrich #MRCF0R030) in 500 µL increments. For each 500 µL increment, each filter was centrifuged at 14,000 rcf for 15-30 min between 500 µL increments to remove SDS-containing buffer. Each filter was then simultaneously washed and reduced using 8M urea, 50 mM ammonium bicarbonate, and 4 mM DTT five times, centrifuging at 14,000 rcf to pass the majority of each wash through the filter between loadings. The filters were then washed using 8 mM iodoacetamide in 50 mM ammonium bicarbonate a total of three times to remove urea and alkylate proteins, followed by 50 mM ammonium bicarbonate two times prior to tryptic digestion. Promega® sequencing-grade trypsin (#MRCF0R030) was then added to each filter in 50 mM ammonium bicarbonate at a 1:50 enzyme-to-protein (w/w) ratio and incubated at 37°C overnight. A secondary tryptic digestion was performed for 4 hours at 37°C the following day using 1:100 w/w trypsin. Resulting peptides were recovered by centrifugation and additional washes of the filter with 50 mM ammonium bicarbonate. Peptides were then dried under vacuum centrifugation and resuspended immediately for LC-MS/MS analysis or phosphopeptide enrichment.

### Phosphopeptide Enrichment

For each phosphopeptide enrichment, 6 mg TiO_2_ (GL Sciences Titansphere, cat #5020-75000) were washed once with 100 µL washing buffer (80% ACN, 15% H2O, 5% TFA). 2 filters of digested peptides (approximately 1.5 mg) were resuspended in binding buffer (1 mL lactic acid, 4 mL washing buffer) for each sample and added to 3 mg TiO_2_ resin. The mixture was allowed to incubate for 30 minutes at RT shaking at 1100 rpm. After binding, the beads were spun down and the supernatant was transferred to another 3 mg TiO_2_ and allowed to bind for an additional 30 minutes. The two TiO2 fractions were then combined and washed using 100 µL of wash buffer with shaking for 10 minutes, repeated a total of 3 times. Phosphopeptides were then eluted twice using 100 µL 15% NH_3_H_2_O, and once using 100 µL 40% ACN, 9% NH_3_H_2_O. The elutions were then combined, dried under vacuum centrifugation, and resuspended for LC-MS/MS analysis.

### NanoLC Separation and nDIA Mass Spectrometry Acquisition

Whole cell lysate and phosphoenriched samples were both analyzed by nano-flow liquid chromatography (Thermo Vanquish Neo) coupled online to a Thermo Orbitrap Astral mass spectrometer. Peptides were loaded directly onto 25 cm Easy Spray Pepmap column (60°C) and eluted at 350 nL/min using a binary solvent system composed of solvent A, 0.1% formic acid in water, and solvent B, 0.1% formic acid in 80% acetonitrile. Unenriched and phosphoenriched samples were separated using 20 min and 40 min acquisitions, respectively, corresponding to 14 and 34 minute gradients starting at 4% A and ending at 35% B.

For nDIA acquisitions, each 3s duty cycle consisted of one full MS1 survey scan followed by a series of DIA MS2 scans. MS1 scans were acquired in the Orbitrap at a resolution of 240,000 over an m/z range of 400–1,000 with AGC target 5e6. MS2 DIA scans were acquired with the Astral detector using fixed 2 Th isolation windows spanned 400-1,000 *m/z*, measuring an m/z range of 150-2000 *m/z* with AGC target 5e4 and max injection time 3 ms. Normalized higher-collision dissociation (HCD) was set to 26%

### DIA-NN Database Search

Raw DIA data were processed using DIA-NN in library-free mode (Demichev et al., 2020). Searches were performed against a predicted spectral library generated from a SwissProt human database supplemented with common contaminants and appended with decoy sequences generated internally by DIA-NN. Enzyme specificity was set to trypsin, allowing up to 2 missed cleavages. Carbamidomethylation of cysteine was specified as a fixed modification, while variable modifications included methionine oxidation and protein N-terminal acetylation. For phosphoenriched samples, phosphorylation of serine, threonine, and tyrosine were included in the search. Precursor charge state was set as 2-4, while precursor m/z range, fragmention m/z range, and other search parameters were set according to the acquisition method. Mass accuracy settings were determined automatically by DIA-NN and retention time prediction and spectral prediction were enabled for predicted library generation. Match-between-runs as well as protein inference were enabled using DIA-NN’s default algorithms. Precursor, peptide, protein group, and phosphosite-level outputs were filtered using a false discovery rate threshold of 1%. For phosphoenrichment analyses, a minimum phosphosite localization confidence of 90% was used in the final analysis

### Quantitative Data Processing

DIA-NN output tables were used for downstream quantitative analysis. Protein group-level, and phosphosite-level quantities were exported for statistical analysis. For global proteome samples, protein group quantities were used to assess protein abundance, sample loading, and reproducibility. For phosphopeptide-enriched samples, phosphopeptide or phosphosite quantities were used for phosphorylation-focused analysis, normalized by protein abundances determined in unenriched samples where possible. Protein-normalized phosphorylation ratios were calculated for treated versus control samples, and significantly regulated phosphosites were defined using combined thresholds for adjusted p-value, fold change, localization confidence, and reproducibility across biological replicates.

### Statistical analyses

Statistical analysis was performed using Prism (Graphpad). Asterisks in figures denote statistical significance as estimated by either Mann–Whitney tests or Chi-square analyses (*, P < 0.05; **, P < 0.01; ***, P < 0.001; ****, P < 0.0001).

## ACKNOWLEDGEMENTS

We thank Daniel I. Martinez for advice on CRISPR-Cas9 experiments, Adeela Syed for advice and assistance with live imaging microscopy, Pauline Nguyen for help with flow cytometry and Helen Hong and Jordan Bond for assistance with quantitative PCR experiments. This study was made possible in part through access to the Optical Biology Core Facility of the Developmental Biology Center, a shared resource supported by the Cancer Center Support Grant (CA-62203) and Center for Complex Biological Systems Support Grant (GM-076516) at the University of California, Irvine P.L.-G. is funded by an NIH R35 grant (GM150786), a Hellman Fellowship and a UCOP Cancer Research Coordinating Committee seed grant (C26CR9985). L.H. is funded by an NIH R35 grant (GM145249). J.V. is funded by the NSF Graduate Research Fellowship Program (NSF-GRFP) (DGE-2235784).

## DATA AVAILABILITY

Raw mass spectrometry data are available via Proteome Xchange with the identifier:

**Project accession:** PXD082726

**Password:** CNoxAVyw5aXX

Original data, cell lines, and plasmids generated in this study are available upon request.

## SUPPLEMENTARY FIGURE LEGENDS

**Figure S1.**
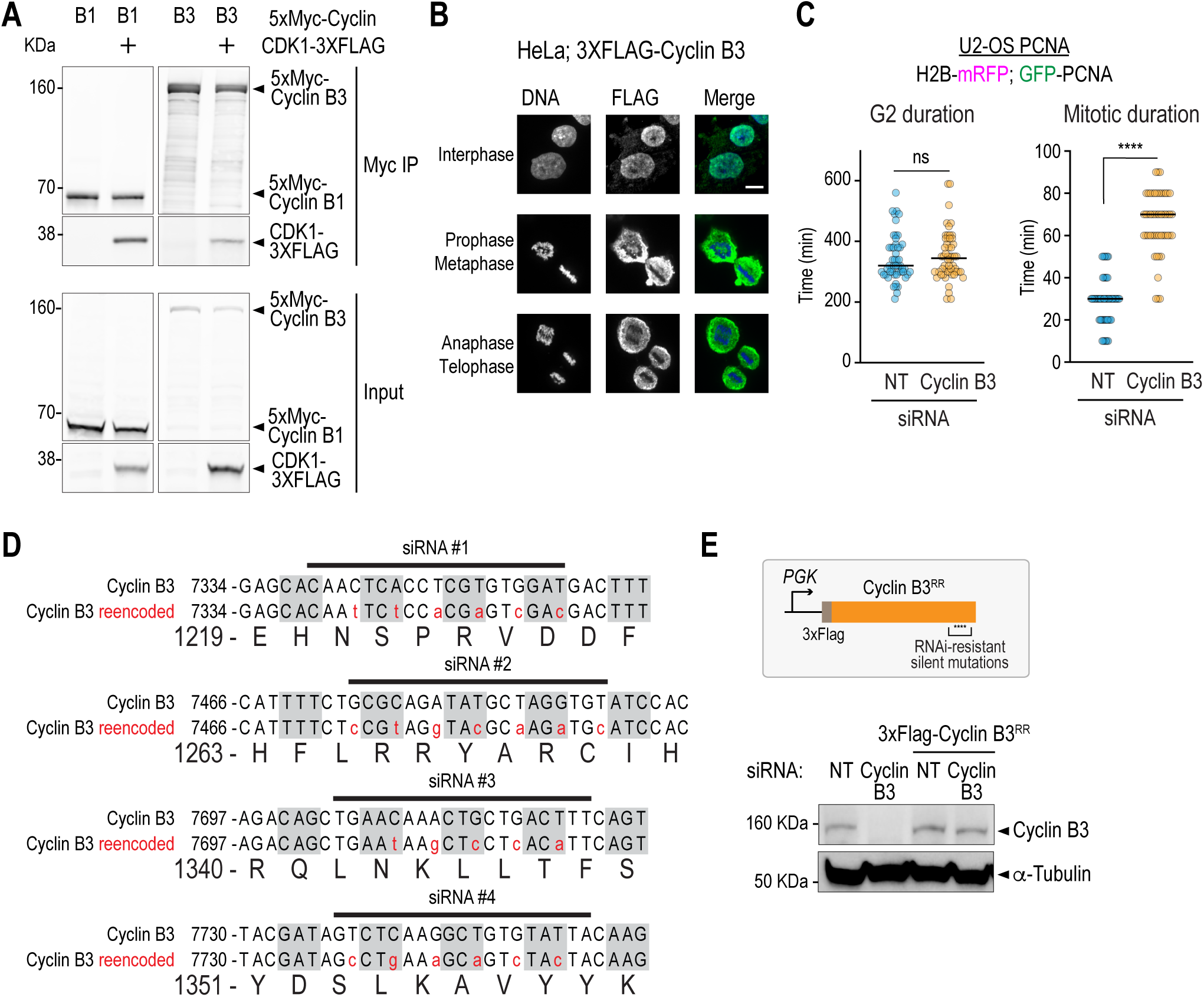
Characterization of cyclin B3 interaction with CDK1 and cell cycle phenotypes. *Related to* Fig. 1 *and* Fig. 2. **(A)** Plasmids encoding for CDK1-3XFLAG were co-transfected into HEK293T cells, along with either 5xMyc-cyclin B1 or 5xMyc-cyclin B3. Cells were subsequently collected and lysed and complexes were pulled down using anti-Myc beads. Note that, under these conditions, 5xMyc-cyclin B1 was more efficient at co-immunoprecipitating CDK1-3XFLAG compared with 5xMyc-cyclin B3, which suggests that cyclin B3 is weaker at interacting with CDK1 compared to cyclin B1. **(B)** Immunofluorescence images of HeLa cells expressing 3xFlag-cyclin B3 and stained with DAPI (blue), α-FLAG (Green) and illustrating examples of cyclin B3 localization in interphase (top), prometaphase and metaphase (middle) and anaphase and telophase (bottom). The images are maximum intensity projections of 4 confocal slices. Note that cyclin B3 localizes to the nucleus in interphase and is largely cytosolic during mitosis. Scale bar, 10 µm. **(C)** Quantification of the duration of G2 *(left)* and mitosis *(right)* phases of the cell cycle from movies of U2-OS cells expressing GFP-PCNA and H2B-mRFP, under the specified conditions. **(D)** Schematics illustrating the binding sites for the cyclin B3 siRNA pool used in this study, as well as the silent mutations engineered in our cyclin B3 reencoded construct (red residues). **(E)** *(top)* Schematic illustrating the construct used for the expression of 3xFlag-tagged, RNAi-resistant (RR) cyclin B3 transgenes. *(bottom)* Immunoblot showing cyclin B3 expression under the indicated conditions. α-Tubulin serves as a loading control. **** represents *P < 0.0001* from Mann-Whitney tests; non-significant (n.s.) is *P > 0.05*.

**Figure S2.**
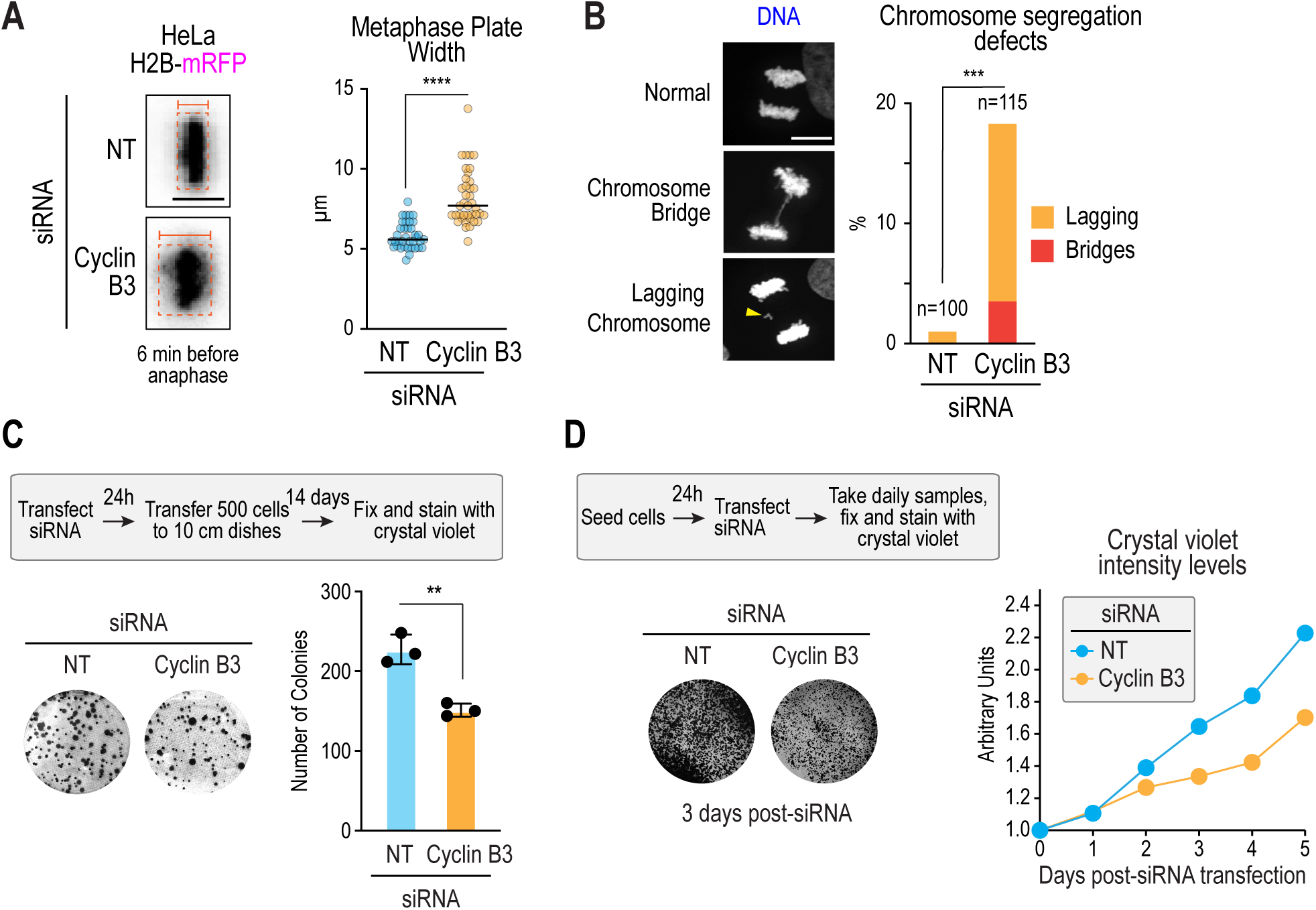
Characterization of cyclin B3 depletion phenotypes. *Related to* Fig. 2. **(A)** *(left)* Representative time-lapse sequences of HeLa cells expressing mRFP-tagged histone H2B and captured 6 minutes before anaphase onset. The red box shows a minimum bounding box encompassing all chromosomes. *(right)* Quantification of metaphase plate width by measuring the width of the minimal bounding box for the specified conditions. Scale bar, 10 µm. **(B)** *(left)* Representative images of anaphase cells stained with DAPI (grey) and showing either chromosome bridges or lagging chromosomes (arrowhead). *(right)* Quantification of chromosome segregation defects for the indicated conditions. Scale bar, 10 µm. **(C)** *(top)* Description of the experimental conditions for the scoring of colony formation. *(bottom left)* Example images of colonies formed by HeLa cells treated with either non-targeting siRNA (NT) or cyclin B3 siRNA. *(bottom right)* quantification of colony numbers under the specified conditions. Error bars are standard deviation. **(D)** *(top left)* Description of the experimental conditions used for the scoring of cell proliferation. *(bottom left)* Example images of cells treated for 3 days with either NT siRNA or cyclin B3 siRNA and stained with crystal violet. *(right)* Growth curves of cells under the specified conditions. For these assays, the crystal violet intensities were scored for each day post-transfection and normalized against values at t=0 days. Each data point represents the average of 6 replicates. Graphs from panels *A* and *C* were analyzed using Mann-Whitney tests; panels *B* was analyzed using Chi square analysis. **** represents *P < 0.0001*; *** represents *P < 0.001*; ** represents *P < 0.01*; non-significant (n.s.) is *P > 0.05*.

**Figure S3.**
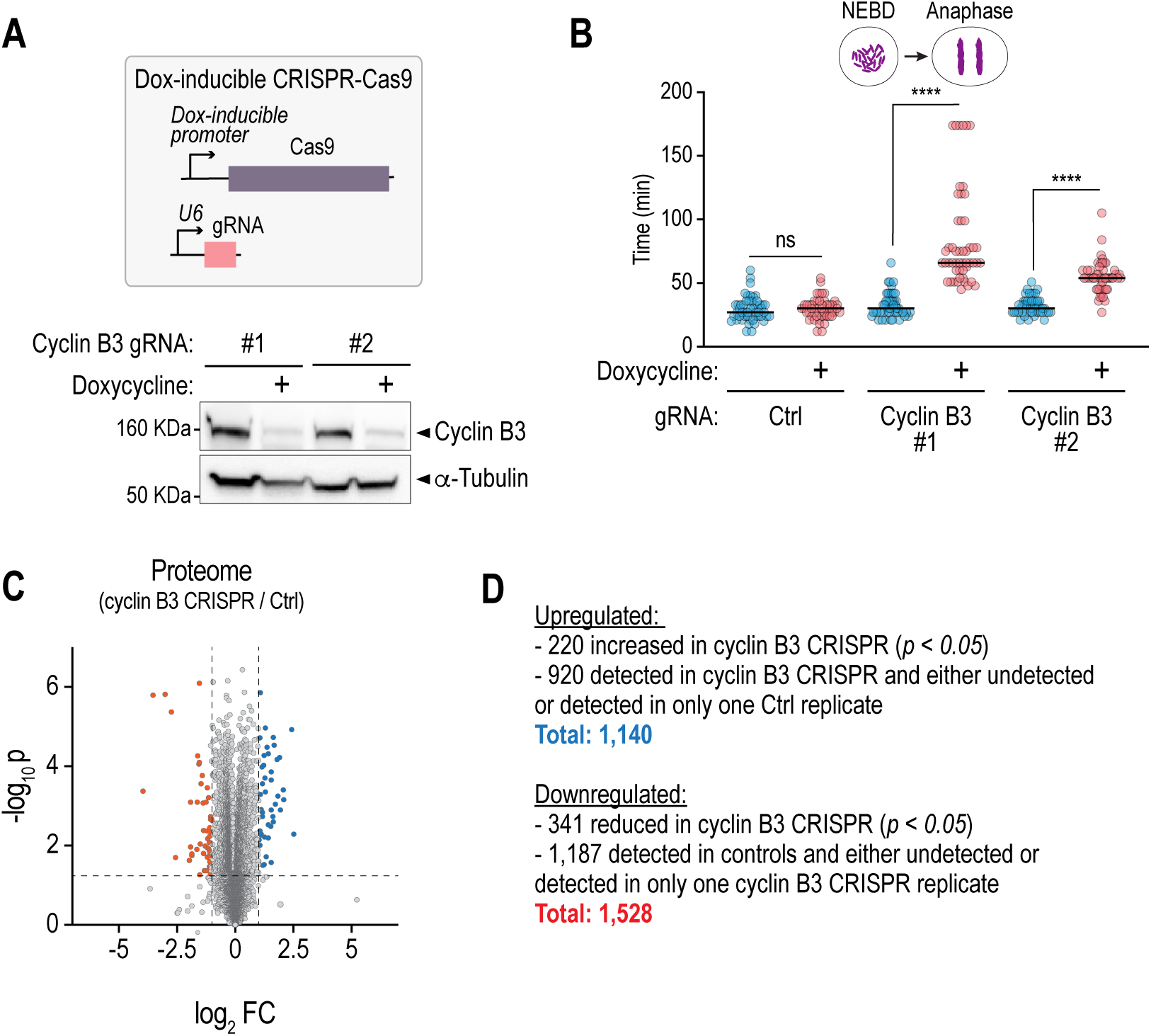
Generation and validation of doxycycline-induced CRISPR-Cas9 cell lines for mass spectrometry analyses. *Related to* Fig. 3. **(A)** *(top)* Schematic illustrating the constructs used for the generation of doxycycline-induced CRISPR-Cas9-mediated editing of cyclin B3. *(bottom)* Immunoblot showing cyclin B3 depletion upon induction of cyclin B3 guide RNAs (gRNA). a-Tubulin serves as a loading control. **(B)** Quantification of the timing from NEBD to Anaphase onset for the specified conditions. **** represents *P < 0.0001* from Mann-Whitney tests; non-significant (n.s.) is *P > 0.05*. **(C)** Volcano plot showing whole proteome changes upon cyclin B3 depletion through CRISPR-Cas9 (gRNA #1). Note that only a small number of proteins either increase or decrease upon cyclin B3 depletion. **(D)** Summary of differentially enriched phosphorylated peptides upon cyclin B3 depletion through CRISPR-Cas9. **** represents *P < 0.0001* from Mann-Whitney tests; non-significant (n.s.) is *P > 0.05*.

**Figure S4.**
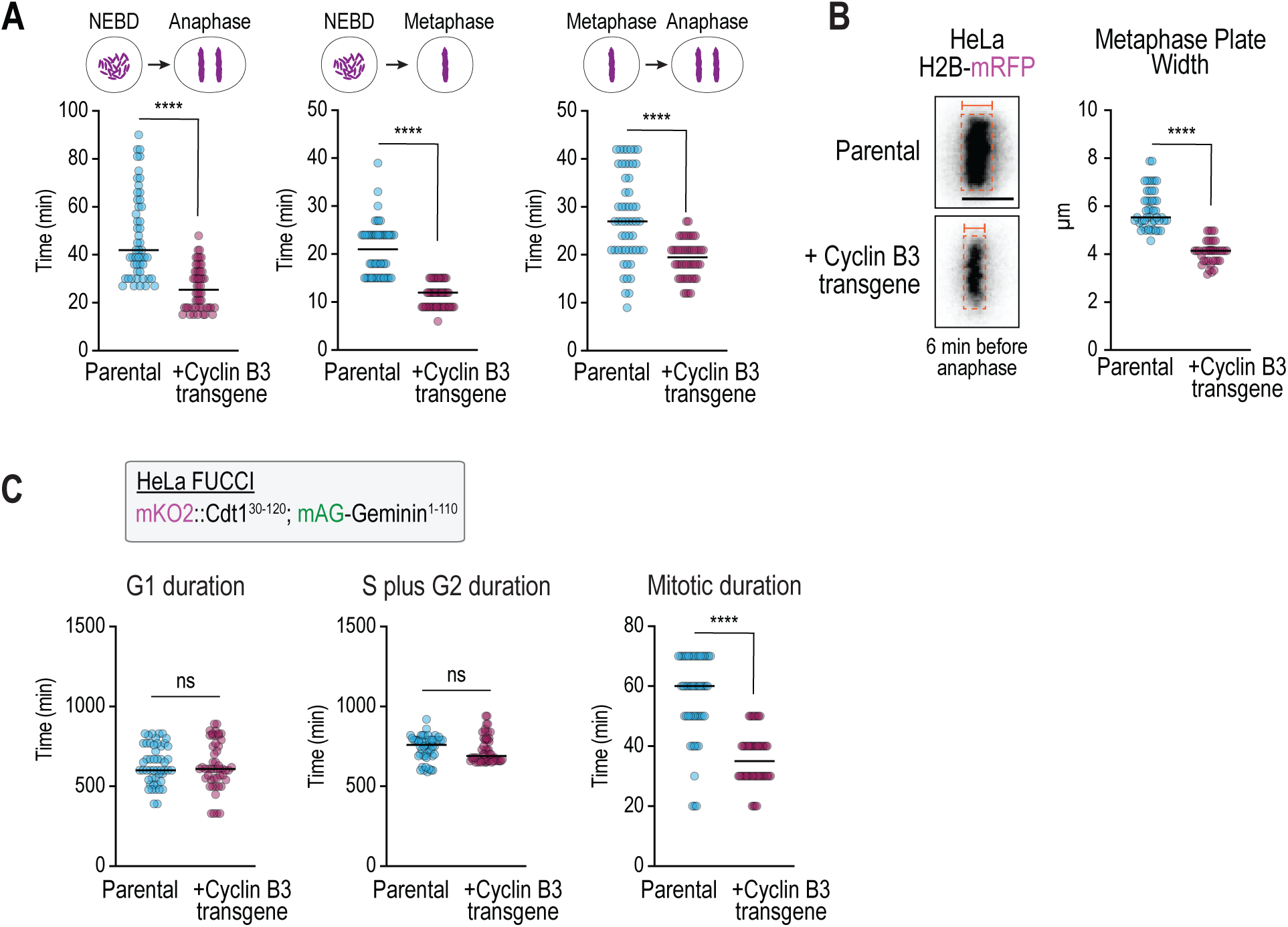
Characterization of the cyclin B3 overexpression phenotype. *Related to* Fig. 4. **(A)** Quantification of the interval from NEBD to Anaphase *(left)*, NEBD to Metaphase *(middle)* and Metaphase to Anaphase *(right)* for the indicated conditions. **(B)** *(left)* Representative time-lapse sequences of HeLa cells expressing mRFP-tagged histone H2B and captured 6 minutes before anaphase onset. The red box shows a minimum bounding box encompassing all chromosomes. Scale bar, 10 µm. *(right)* Quantification of metaphase plate width by measuring the width of the minimal bounding box for the specified conditions. **(C)** Quantification of the duration of G1, S/G2 and mitosis in HeLa cells expressing the FUCCI marker. **** represents *P < 0.0001* from Mann-Whitney tests;; non-significant (n.s.) is *P > 0.05*.

**Figure S5.**
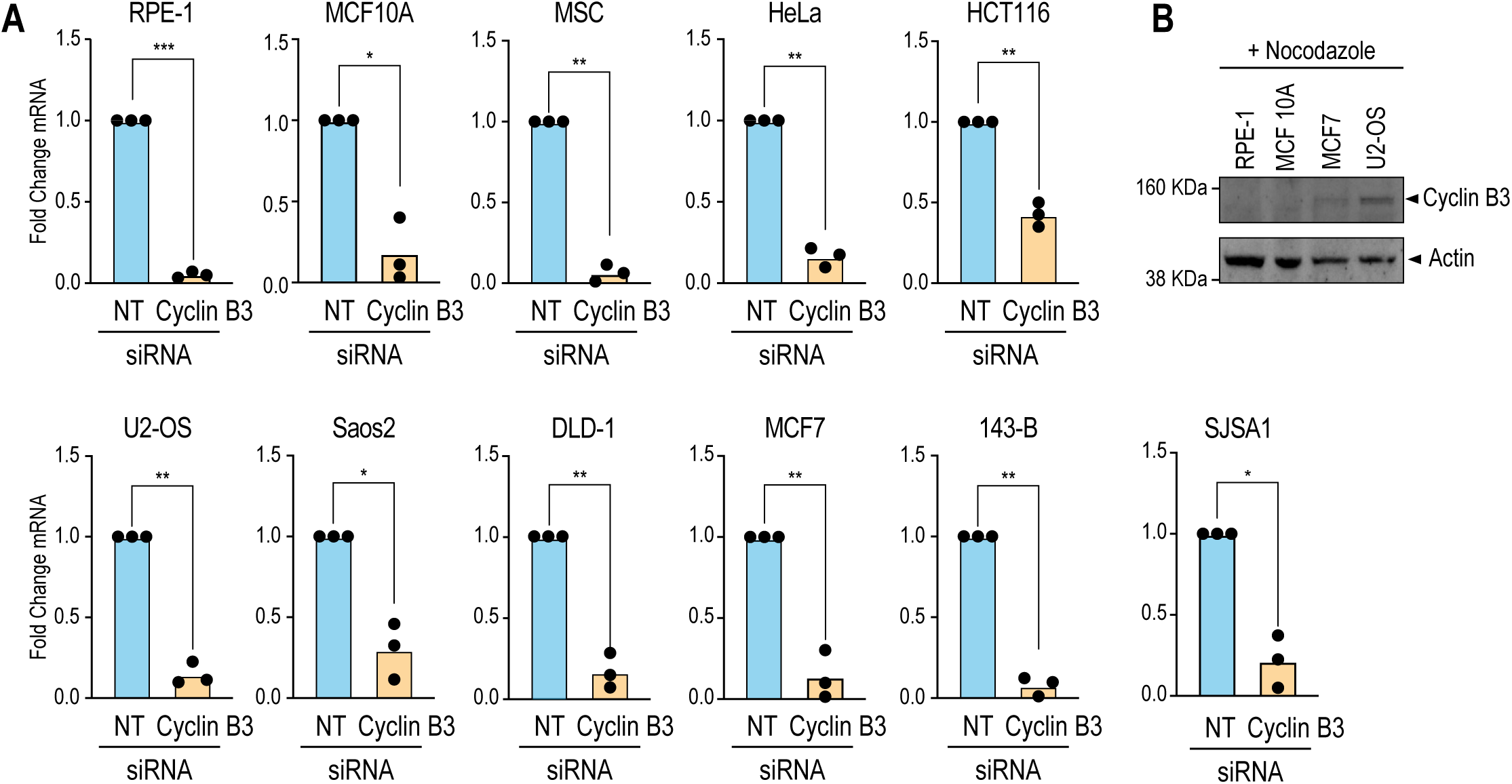
Expression of cyclin B3 in cancer cell lines and validation of cyclin B3 depletion through quantitative PCR (qPCR). *Related to* Fig. 4. (A) Quantification of cyclin B3 mRNA levels in cells treated with either NT siRNA or cyclin B3 siRNA. Note that all tested cell lines showed efficient cyclin B3 depletion at the mRNA level. **(B)** Immunoblot showing cyclin B3 expression in mitotically arrested RPE-1, MCF 10A, MCF7 and U2-OS cells. Actin serves as a loading control. **** represents *P < 0.0001* from Mann-Whitney tests; *** represents P *< 0.001*; ** represents *P < 0.01*; * represents *P < 0.05*; non-significant (n.s.) is *P > 0.05*.

